# Improved CRISPR/Cas9 Off-target Prediction with DNABERT and Epigenetic Features

**DOI:** 10.1101/2025.04.16.649101

**Authors:** Kai Kimata, Kenji Satou

## Abstract

CRISPR/Cas9 genome editing is a powerful tool in genetic engineering and gene therapy; however, off-target effects pose significant challenges for clinical applications. Accurate prediction of these unintended edits is crucial for ensuring safety and efficacy. In this study, we propose a novel approach that integrates DNABERT, a pre-trained DNA language model, with epigenetic features to improve off-target effect prediction. We evaluated DNABERT-based models against five state-of-the-art baseline models (GRU-Emb, CRISPR-BERT, CRISPR-HW, CRISPR-DIPOFF, and CrisprBERT) using four key performance metrics (F1-score, MCC, ROC-AUC, and PR-AUC). Additionally, we conducted ablation studies to assess the impact of DNABERT’s pre-training and epigenetic features, demonstrating that both significantly enhance predictive performance. Furthermore, we explored an ensemble modeling approach, which achieves superior prediction accuracy compared to individual models. Finally, we visualized DNABERT’s attention weights to gain insights into its decision-making process, revealing biologically relevant patterns in off-target recognition. The source codes used in this study are available at github.com/kimatakai/CRISPR_DNABERT.

## Introduction

CRISPR(Clustered Regularly Interspaced Short Palindromic Repeats)/Cas9 genome editing technology, co-invented by Emmanuelle Charpentier and Jennifer Doudna in 2012, has been widely used in various fields such as genetic engineering and gene therapy research [1–4]. This system utilizes a single-guide RNA (sgRNA) to direct the Cas9 nuclease to a target DNA site, resulting in DNA cleavage [5, 6]. The first 20 nucleotides of the sgRNA are complementary to the target DNA sequence, followed by a sequence known as the protospacer adjacent motif (PAM), typically ‘NGG’ [7].

The target specificity of Cas9 is governed by the 20-nt sequence of the sgRNA and the PAM sequence adjacent to the target sequence in the genome. However, in practice, off-target effects may occur, leading to DNA cleavage even when there are several base pair mismatches between the sgRNA and the genomic sequence [7–9].

Editing of non-target DNA due to off-target effects can potentially disrupt normal gene function. This poses a significant problem for the clinical application of CRISPR/Cas9 genome editing [10–12]. Therefore, predicting the DNA regions where off-target effects may occur is crucial for understanding their potential impacts in advance.

In recent years, various computational methods have been developed to predict off-target effects [13, 14]. Many deep learning-based prediction models have been developed recently, demonstrating superior performance compared to traditional empirical-based methods that rely on prior knowledge and a series of algorithms [10, 13–15]. In particular, attention-based deep learning models leveraging the BERT architecture, widely used in natural language processing, have been introduced for CRISPR/Cas9 off-target prediction [16–23]. However, most of the currently available deep learning models for CRISPR/Cas9 off-target prediction have been trained exclusively on off-target effect datasets, and no studies have explored the use of large-scale, pre-trained models trained on genomic corpora such as the human genome.

While the sequence complementarity between sgRNA and target DNA is a key determinant of Cas9 binding and cleavage activity, accumulating evidence suggests that epigenetic modifications also influence CRISPR/Cas9 activity [14, 24]. Several studies have incorporated epigenetic features into deep learning models to improve off-target effect prediction [20, 25–27]. However, most CRISPR/Cas9 off-target prediction research remains focused solely on sequence-based information, overlooking the potential impact of epigenetic factors.

In this study, we proposed a novel approach that integrates DNABERT, a pre-trained model designed for DNA sequences, into CRISPR/Cas9 off-target effect prediction [28]. Furthermore, we incorporated epigenetic features to enhance predictive performance. To evaluate the effectiveness of our approach, we compared it against five state-of-the-art models (GRU-Emb, CRISPR-BERT, CRISPR-HW, CRISPR-DIPOFF, and CrisprBERT). We ensured the robustness of our results by conducting cross-validation with multiple random seeds and evaluating performance using four key metrics: F1-score, MCC, ROC-AUC, and PR-AUC. Additionally, to assess the impact of DNABERT’s pre-training and the inclusion of epigenetic features, we performed two ablation studies. Finally, we visualized the attention weights of DNABERT to gain interpretability insights into its decision-making process.

The source codes used in this study are available at: github.com/kimatakai/CRISPR_DNABERT

## Materials and Methods

### Datasets

In this study, we used high-throughput CRISPR/Cas9 off-target datasets from CHANGE-seq (in vitro) and GUIDE-seq (in cellula), which have been curated to include off-target sites (OTS) with bulges, to train the model and compare the proposed method with other deep learning-based approaches [24, 29, 30].

CHANGE-seq is a method for high-precision and scalable measurement of genome-wide CRISPR/Cas9 nuclease activity. This technique, based on tagmentation (DNA fragmentation and ligation), enables efficient genome-wide analysis of Cas9 activity in vitro, generating massive datasets. The CHANGE-seq dataset consists of 202,040 active OTS and 4,734,238 inactive OTS associated with 110 distinct sgRNAs.

GUIDE-seq is an unbiased method for detecting double-strand breaks (DSBs) introduced by CRISPR RNA-guided nucleases in live cells. The GUIDE-seq dataset is a subset of the CHANGE-seq dataset, covering 58 sgRNAs and consisting of 1,520 active OTS and 2,118,820 inactive OTS.

We used these datasets in three experimental setups:

Scenario 1: Training and testing on the CHANGE-seq dataset.
Scenario 2: Training and testing on the GUIDE-seq dataset.
Scenario 3: Training on the CHANGE-seq dataset, followed by additional training on the GUIDE-seq dataset, and testing on the GUIDE-seq dataset.

### Epigenetic information and off-target effect

GUIDE-seq detects DSBs in living cells, and the detected OTS are influenced by chromatin accessibility, other epigenetic factors, and cellular conditions [30, 31]. Prior studies indicate that GUIDE-seq OTS are more frequently observed in highly expressed genes than CHANGE-seq OTS and are associated with open chromatin (ATAC-seq), active promoters (H3K4me3), and enhancers (H3K27ac) (Fig 1A) [24]. These findings suggest that GUIDE-seq results are influenced by chromatin state in living cells.

**Fig 1.**
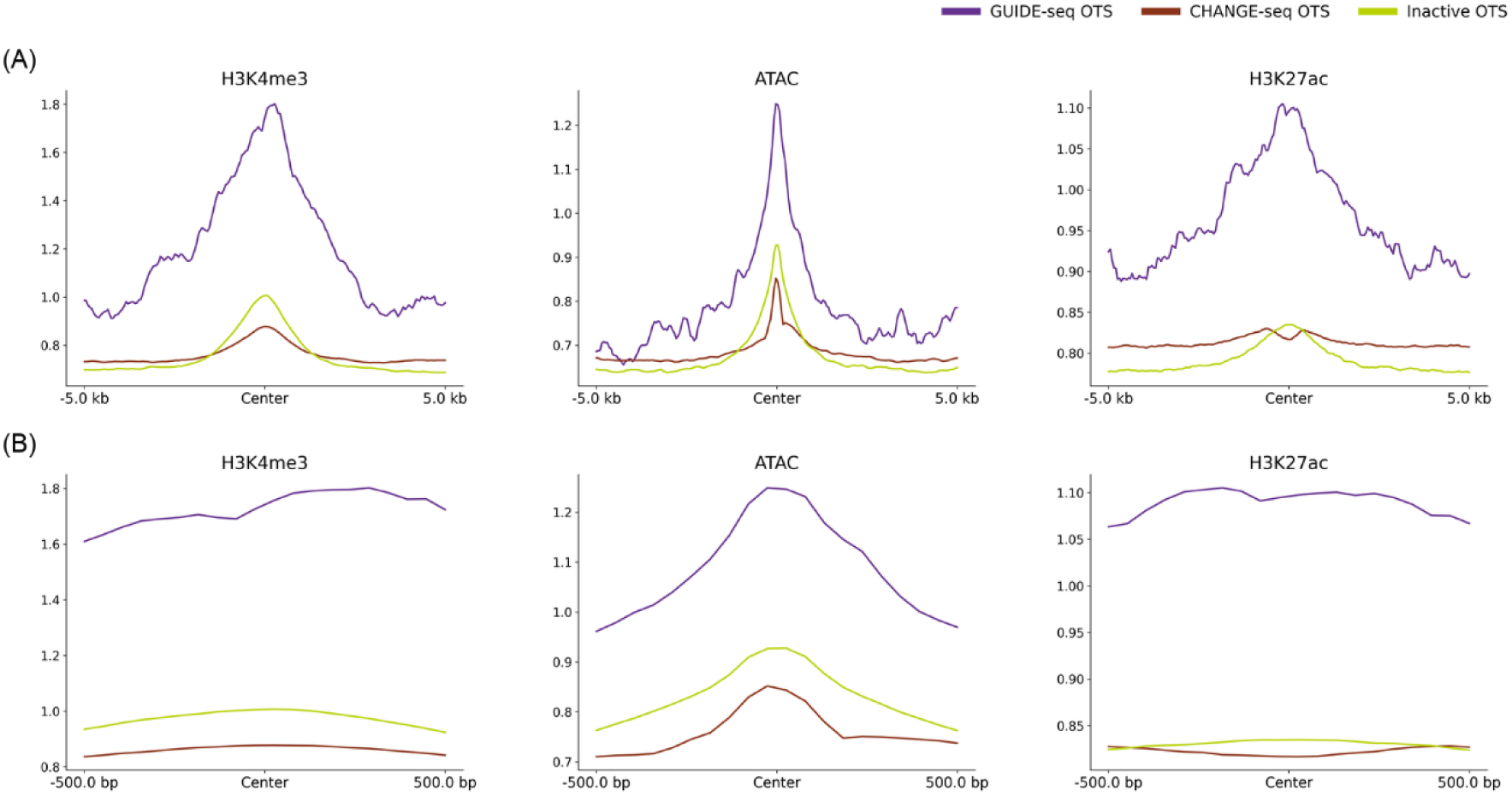
Comparison of epigenetic signals at OTS in CRISPR/Cas9: Visualization of epigenetic markers detected by GUIDE-seq and CHANGE-seq. This figure shows line graph of the average epigenetic signa level at OTS detected by GUIDE-seq and CHANGE-seq, as well as inactive OTS where no off-target effects occurred. The epigenetic quantities represented are the average signal values for H3K4me3, ATAC-seq, and H3K27ac. Panels (A) and (B) display the signal values within ±5.0kb and ±500bp window centered on the OTS, respectively.

This difference likely results from CHANGE-seq, like CIRCLE-seq, using purified genomic DNA, which enables the detection of OTS without chromatin structure constraints [32, 33].

Based on these findings, we hypothesized that incorporating epigenetic information could improve predictive accuracy, particularly when using the GUIDE-seq dataset (Scenarios 2 and 3). Therefore, we integrated H3K4me3, ATAC-seq, and H3K27ac data into the model to capture chromatin accessibility and regulatory element activity. Epigenetic data (H3K4me3, ATAC-seq, and H3K27ac) were obtained from the Gene Expression Omnibus (GSE149363).

### Cross validation strategy

To evaluate predictive performance, we employed 10-fold cross-validation following the method of Yaish et al. [30, 34]. In this approach, the sgRNAs were divided into 10 subsets, with each iteration using one subset as the test set and the remaining subsets as the training set.

For the CHANGE-seq dataset, performance was evaluated for each sgRNA in each iteration of the cross-validation. For the GUIDE-seq dataset, performance was assessed on the entire subset in each iteration.

In the transfer learning scenario, previous studies ensured that sgRNAs in the training subsets of the CHANGE-seq dataset did not overlap with those in the test subsets of the GUIDE-seq dataset. This design prevented information leakage and enabled a fair evaluation of the model’s ability to generalize across datasets.

### Data preprocessing

In accordance with previous studies, active OTS experimentally identified with read counts below 100 were labeled as positive class [30].

Both the CHANGE-seq and GUIDE-seq datasets exhibited extreme class imbalance, with ratios of approximately 23:1 and 1394:1, respectively. Given that the batch sizes of the deep learning models used in this study ranged from 64 to 512, batches without any positive class samples could occur, particularly in the GUIDE-seq dataset. This could potentially lead to overfitting.

To mitigate this, random downsampling of the negative class in the training data was performed, reducing it to 20% of its original size to maintain class balance within each batch. A fixed random seed was applied to ensure that all models used the same downsampled training dataset. The test dataset remained unaltered throughout the process.

For each OTS, epigenetic data for H3K4me3, ATAC-seq, and H3K27ac were processed through several steps. First, a 1000 bp region centered around the first base of the cleavage site was extracted, covering 500 bp upstream and downstream (Fig 1B). Next, the signal within this region was averaged into 50 bp bins, resulting in a 20-dimensional vector. Additionally, the mean value for each feature was calculated and appended to the vector, expanding the vector to 21 dimensions. Finally, the processed vectors for H3K4me3, ATAC, and H3K27ac were concatenated, yielding a 63-dimensional feature vector.

### Repeated experiments

To account for the potential influence of randomness, such as random sampling and the initialization of deep learning model parameters, 10-fold cross-validation was conducted five times with different random seeds. This repetition helped ensure the robustness and reliability of the results.

### DNABERT fine-tuning

DNABERT is a model that extends the BERT architecture — originally developed for natural language processing — to DNA sequences [28]. In this study, we used the 3-mer DNABERT model, which had been pre-trained on a masked language model (MLM) task. The pre-trained model was obtained from Hugging Face. We implemented our approach using transformers v4.48.3 and pytorch v2.5.1.

To adapt DNABERT for off-target prediction, we fine-tuned it in two stages (Fig 2A). First, we trained the model on a mismatch position prediction task, where the objective was to predict whether a mismatch occurred at each position in a given DNA-sgRNA sequence pair. This task was designed to help DNABERT learn the pairing relationship between the two sequences. For this task, we generated random DNA-sgRNA sequence pairs for training. We used a batch size of 8, a learning rate of 0.00002, and trained the model for five epochs during this stage.

**Fig 2.**
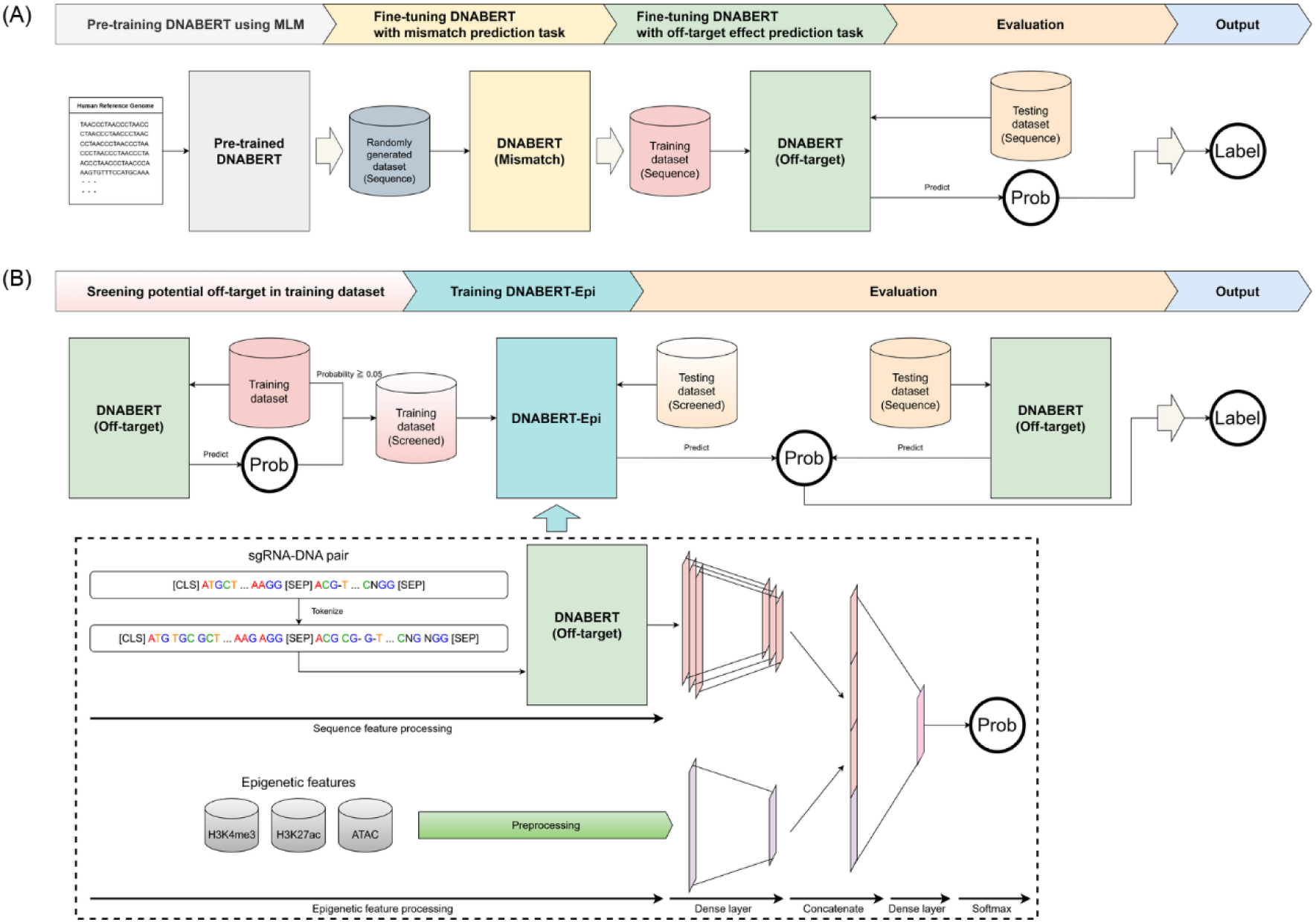
Overview of DNABERT fine-tuning and DNABERT-Epi for off-target effect prediction. (A) DNABERT Fine-tuning: The DNABERT model, pre-trained on a masked language model (MLM) task, is fine-tuned in two stages. The first stage involves training on a mismatch position prediction task, where the model learns to recognize mismatches in DNA-sgRNA sequence pairs. In the second stage, the model is further fine-tuned for binary classification to predict off-target effects. The vocabulary is expanded to include 3-mer tokens with a bulge character (“-”), and input sequences are tokenized in the format: [CLS], DNA 3-mer tokens, [SEP], sgRNA 3-mer tokens, [SEP]. (B) DNABERT-Epi model: DNABERT-Epi integrates epigenetic features to enhance off-target effect prediction. The fine-tuned DNABERT model first estimates off-target probabilities. Samples with predicted probabilities ≧ 0.05 are processed further by DNABERT-Epi, while samples with probabilities < 0.05 retain DNABERT’s output as the final prediction. DNABERT-Epi combines sequence-based embeddings from DNABERT with epigenetic features through a feedforward neural network (FNN). The final probability is computed by averaging DNABERT-Epi and DNABERT outputs for high-probability samples, while low-probability samples use DNABERT’s output directly.

Next, we fine-tuned the DNABERT model, which had been trained on the mismatch prediction task, for binary classification to predict off-target effects. In this second fine-tuning, we used a batch size of 256, a learning rate of 0.00002, and trained the model for five epochs.

Before fine-tuning, we modified DNABERT’s vocabulary by adding 3-mer tokens that included the bulge character (“-”). Input sequences were tokenized in the following format:

[CLS], DNA 3-mer tokens, [SEP], sgRNA 3-mer tokens, [SEP]

This two-stage fine-tuning approach aimed to leverage DNABERT’s pre-training while enabling it to effectively model off-target interactions.

### Incorporating epigenetic features into off-target prediction

To enhance the accuracy of off-target effect prediction, we propose DNABERT-Epi, an extension of the fine-tuned DNABERT model that incorporates epigenetic features (Fig 2B). The prediction process consists of two stages. First, the fine-tuned DNABERT model estimates the off-target probability for each sample in the training dataset. Samples predicted with a probability of 0.05 or higher are treated as potential positives and selected for further training in DNABERT-Epi.

In the DNABERT-Epi model, the tokenized sequence and epigenetic features are first fed as inputs. The tokenized sequence is processed by the DNABERT unit to extract three types of embeddings: the CLS token embedding, the maximum-pooled representation of all token embeddings, and the mean-pooled representation of all token embeddings. Each of these embeddings is processed through a feedforward neural network (FNN) layer, followed by a ReLU activation function and a dropout layer with a dropout rate of 0.1, resulting in a 32-dimensional vector. Simultaneously, the preprocessed epigenetic features are also passed through an FNN layer, a ReLU activation function, and a dropout layer with a dropout rate of 0.1, transforming them into a 32-dimensional vector. These four feature sets are then concatenated to form a 128-dimensional feature representation. This representation is further processed through a 32-dimensional FNN layer, a ReLU activation function, and a dropout layer before being passed to the final output layer and a Softmax layer, which computes the probability of off-target effects.

For training DNABERT-Epi, we used a batch size of 256, a learning rate of 0.0001, and trained the model for five epochs.

During inference, the same screening process is applied. Samples predicted by the fine-tuned DNABERT model with a probability of 0.05 or higher are input into the trained DNABERT-Epi model, and the final output probability is obtained by averaging the predictions from the DNABERT-Epi model and the DNABERT model. For samples with predicted probabilities below 0.05, the DNABERT model’s output is used directly as the final prediction.

### Baseline models and ensemble approach

To evaluate the performance of DNABERT and DNABERT-Epi, we compared against five state-of-the-art deep learning-based models: GRU-Emb, CRISPR-BERT, CRISPR-HW, CRISPR-DIPOFF, and CrisprBERT [22, 23, 30, 35, 36]. These models were selected as they have demonstrated high performance in CRISPR off-target prediction. Since publicly available implementations were not always consistent due to variations in the types and versions of libraries used, we reimplemented these models in PyTorch v2.5.1 based on their original papers and source codes to ensure fair comparisons under identical experimental conditions. To enhance reproducibility and facilitate further research, we have made our reimplementation publicly available on GitHub.

Some modifications were necessary to ensure compatibility with our dataset. Specifically, CRISPR-DIPOFF and CrisprBERT did not originally support bulge-containing input sequences, so we modified their data processing components accordingly. Although we initially considered CRISPR-M, the specific model used in the original study could not be clearly identified among several implementations available on GitHub [20]. Consequently, we excluded it from our comparisons.

For hyperparameter settings, we primarily followed the values reported in the respective papers and source codes (Table 1). However, for CrisprBERT, the default number of training epochs in the source code was set to 400. To reduce computational cost and mitigate overfitting, we adjusted it to 10 epochs. Additionally, as smaller epochs are generally preferable for preventing overfitting during transfer learning, we reduced the number of epochs by half when fine-tuning on the GUIDE-seq dataset.

**Table 1.** Hyperparameters of each baseline model.

| Model name | Epochs | Batch size | Learning rate |
| --- | --- | --- | --- |
| GRU-Emb | 10 | 512 | 0.005 |
| CRISPR-BERT | 30 | 256 | 0.0001 |
| CRISPR-HW | 30 | 128 | 0.003 |
| CRISPR-DIPOFF | 50 | 64 | 0.0001 |
| CrisprBERT | 10 | 128 | 0.00002 |

To further enhance prediction robustness, we developed an ensemble approach that integrates all baseline models along with DNABERT and DNABERT-Epi. Specifically, we employed a soft voting strategy, in which the final prediction was determined by averaging the probability scores output by each model. This approach aimed to leverage the complementary strengths of different architectures, leading to more stable and accurate predictions.

### Evaluation metrics

In this study, we selected F1 score (F1), Matthews correlation coefficient (MCC), receiver operating characteristic-area under the curve (ROC-AUC), and precision-recall area under the curve (PR-AUC) as evaluation metrics, considering the imbalance in the dataset used in this study.

The F1 score is the harmonic mean of precision and recall, making it suitable for assessing the overall classification performance in imbalanced datasets. The MCC accounts for all four elements of the confusion matrix and provides a balanced assessment of classification performance, even for imbalanced data. It ranges from -1 to 1, with values closer to 1 indicating better classification performance. ROC-AUC evaluates the trade-off between the true positive rate (TPR) and the false positive rate (FPR) across different thresholds, indicating the model’s overall discriminative ability. PR-AUC measures the trade-off between precision and recall, offering a more informative assessment of model performance in imbalanced datasets. By employing these metrics, we ensure a comprehensive evaluation of classification performance, particularly in the presence of data imbalance.

### Evaluation of pre-training effectiveness

To evaluate the impact of DNABERT’s pre-training on CRISPR/Cas9 off-target prediction, we conducted an ablation study. This study compared the performance of a pre-trained DNABERT model with a randomly initialized DNABERT model, where both models were fine-tuned under identical conditions. The objective was to investigate whether pre-training plays a significant role in improving model performance.

In the training from scratch condition, we initialized the DNABERT model with random weights, rather than using pretrained weights. The model was then fine-tuned through two sequential steps: first, the model underwent fine-tuning on a mismatch prediction task using paired sequences, following the same procedure as with the pretrained DNABERT. Afterward, the model was fine-tuned on the off-target effect prediction task. This approach allowed us to directly compare the pretrained model’s effectiveness with a model that was trained from scratch.

The architecture of both the pretrained DNABERT and the randomly initialized DNABERT remained identical, as both utilized the same transformer-based model. All hyperparameters, including the learning rate, batch size, and optimizer settings, were held constant to ensure that the comparison between the models was fair.

### Evaluation of the impact of epigenetic features

To evaluate the effect of incorporating epigenetic features into the off-target prediction model, we performed an ablation study comparing the DNABERT model with the extended DNABERT-Epi model, which includes epigenetic features. The goal was to determine whether the performance improvements observed in DNABERT-Epi were due to the inclusion of epigenetic information or the screening process that identifies potential positive samples for further training.

In the DNABERT-Epi model, the prediction process was divided into two stages. First, the fine-tuned DNABERT model was used to estimate the off-target probability for each sample in the training dataset. Samples with a predicted probability of 0.05 or higher were identified as potential positives and selected for further training in the DNABERT-Epi model, which incorporates both sequence data and epigenetic features. For the ablation study, we compared the performance of DNABERT-Epi, which includes epigenetic features, to a version of the model that uses the same screening procedure but excludes epigenetic features.

In the ablation version, only the tokenized DNA sequence was used as input to the model, with the epigenetic features removed from the input layer. The sequence data was processed through the same transformer architecture and pooling layers as DNABERT-Epi, but without the epigenetic feature processing. The screening step, which selects samples with a predicted probability of 0.05 or higher, was applied to both models in the same way. Both models were trained using the same hyperparameters. This allowed us to assess whether the observed performance improvement in DNABERT-Epi was primarily due to the inclusion of epigenetic features or the screening process itself.

### Attention weights visualization

Transformer-based NLP models, such as BERT and GPT-3, employ a multi-head attention mechanism. By visualizing the attention weights, it is possible to understand how the model attends to different positions in the input sequence, which can help interpret the model’s decision-making process [37, 38]. The DNABERT model used in this study consists of 12 transformer layers, each utilizing a multi-head self-attention mechanism with 12 attention heads. We visualized the attention weights of each transformer layer by treating them as connections between tokens. The attention weights for each layer were obtained by averaging the values across the 12 attention heads. In this study, we specifically visualized the differences in attention weights between active and inactive OTS.

## Results

### Scenario 1 results

The performance of each model was evaluated using the CHANGE-seq dataset (S1 Fig, S1 and S2 Table). DNABERT achieved the highest ROC-AUC (0.9187), significantly outperforming other models (p-value < 0.01). For PR-AUC, CRISPR-BERT and CrisprBERT exhibited the highest scores, while F1 and MCC were comparable among GRU-Emb, CRISPR-BERT, and CrisprBERT, with no significant differences. The ensemble model significantly outperformed all other models across all key metrics. Additional evaluation metrics, including Accuracy, Recall, and Precision, are provided in the supporting information (S2 Fig and S1 and S2 Table).

### Scenario 2 results

The evaluation results on the GUIDE-seq dataset are summarized in the supporting information (S3 Fig and S3 and S4 Table). DNABERT-Epi and CRISPR-BERT achieved the highest performance in terms of F1 and MCC, with no statistically significant difference between them. However, DNABERT-Epi significantly outperformed DNABERT in both metrics. In terms of ROC-AUC, GRU-Emb, CRISPR-BERT, CrisprBERT, and DNABERT-Epi demonstrated high performance with no significant differences among them, whereas DNABERT-Epi was significantly better than DNABERT. Regarding PR-AUC, DNABERT-Epi achieved the highest score, significantly outperforming all other models. Moreover, the ensemble model exhibited statistically significant superiority across these evaluation metrics. Detailed results for additional metrics can be found in the supporting information (S4 Fig and S3 and S4 Table).

### Scenario 3 results

The evaluation results after pre-training on the CHANGE-seq dataset and additional training on the GUIDE-seq dataset are presented in the figures and the supporting information (Fig 2, S5 Fig, and S5 and S6 Table). DNABERT-Epi achieved significantly higher performance than other models in terms of F1 and MCC. For ROC-AUC, CrisprBERT exhibited the highest score, showing a statistically significant difference compared to other models. In terms of PR-AUC, both CrisprBERT and DNABERT-Epi demonstrated strong performance, although no significant difference was observed between these two models. Furthermore, the ensemble model outperformed all other models across all evaluation metrics with statistical significance. Detailed results for additional metrics and statistical analyses can be found in the supporting information (S5 Fig and S5 and S6 Table).

### Results of ablation study on pre-training effectiveness

The ablation study aimed to assess the importance of DNABERT’s pretraining by comparing the performance of the pre-trained model with that of a randomly initialized model trained from scratch on the GUIDE-seq dataset (Scenario 3). The results of this comparison are summarized in Table 2. Clearly, the randomly initialized model was unable to distinguish between active and inactive OTS, suggesting that it failed to learn meaningful patterns for this task.

**Table 2.** Performance comparison of DNABERT with pre-trained and from scratch.

|  | **F1** | **MCC** | **ROC-AUC** | **PR-AUC** |
| --- | --- | --- | --- | --- |
| Pre-trained DNABERT | $0.4152 \pm 0.0928$ | $0.4312 \pm 0.0766$ | $0.9809 \pm 0.0118$ | $0.4554 \pm 0.0894$ |
| DNABERT from scratch | $0.0000 \pm 0.0000$ | $0.0000 \pm 0.0000$ | $0.5182 \pm 0.0768$ | $0.0076 \pm 0.0170$ |
| p-value | $3.778 \times 10^{-10}$ | $3.778 \times 10^{-10}$ | $8.882 \times 10^{-16}$ | $8.882 \times 10^{-16}$ |

Additionally, the training loss curves during the mismatch prediction fine-tuning phase further emphasize the benefit of pre-training (S6 Fig). The pre-trained model exhibited a consistent decrease in loss as training progressed, while the model trained from scratch showed a flat loss curve, indicating a lack of meaningful learning.

### Results of ablation study on the effect of epigenetic features

To evaluate whether the inclusion of epigenetic features contributes to improved performance in off-target prediction, we compared DNABERT-Epi with and without epigenetic features on the GUIDE-seq dataset (Scenario 3). The results of this ablation study are summarized in Table 3.

**Table 3.** Performance comparison of DNABERT-Epi with and without epigenetic features.

|  | **F1** | **MCC** | **ROC-AUC** | **PR-AUC** |
| --- | --- | --- | --- | --- |
| DNABERT-Epi with epigenetic | $0.4302 \pm 0.0968$ | $0.4450 \pm 0.0781$ | $0.9809 \pm 0.0118$ | $0.4612 \pm 0.0835$ |
| DNABERT-Epi without epigenetic | $0.4257 \pm 0.0930$ | $0.4396 \pm 0.0776$ | $0.9809 \pm 0.0118$ | $0.4579 \pm 0.0843$ |
| p-value | 0.0247 | 0.0096 | 0.0362 | 0.0131 |

As shown in Table 3, DNABERT-Epi with epigenetic features outperformed the model without epigenetic AUC were relatively small, statistical analysis revealed that these differences were significant (p < 0.05 for all metrics).

### Attention weights visualization of DNABERT

To gain insight into how DNABERT distinguishes between active and inactive OTS, we visualized the attention weights at each transformer layer. The visualization was generated by averaging the attention weights across all attention heads and then computing the difference between active and inactive OTS. The results are presented in Fig 4, where red lines indicate higher attention weights inactive OTS, and blue lines indicate higher attention weights in inactive OTS.

**Fig 3.**
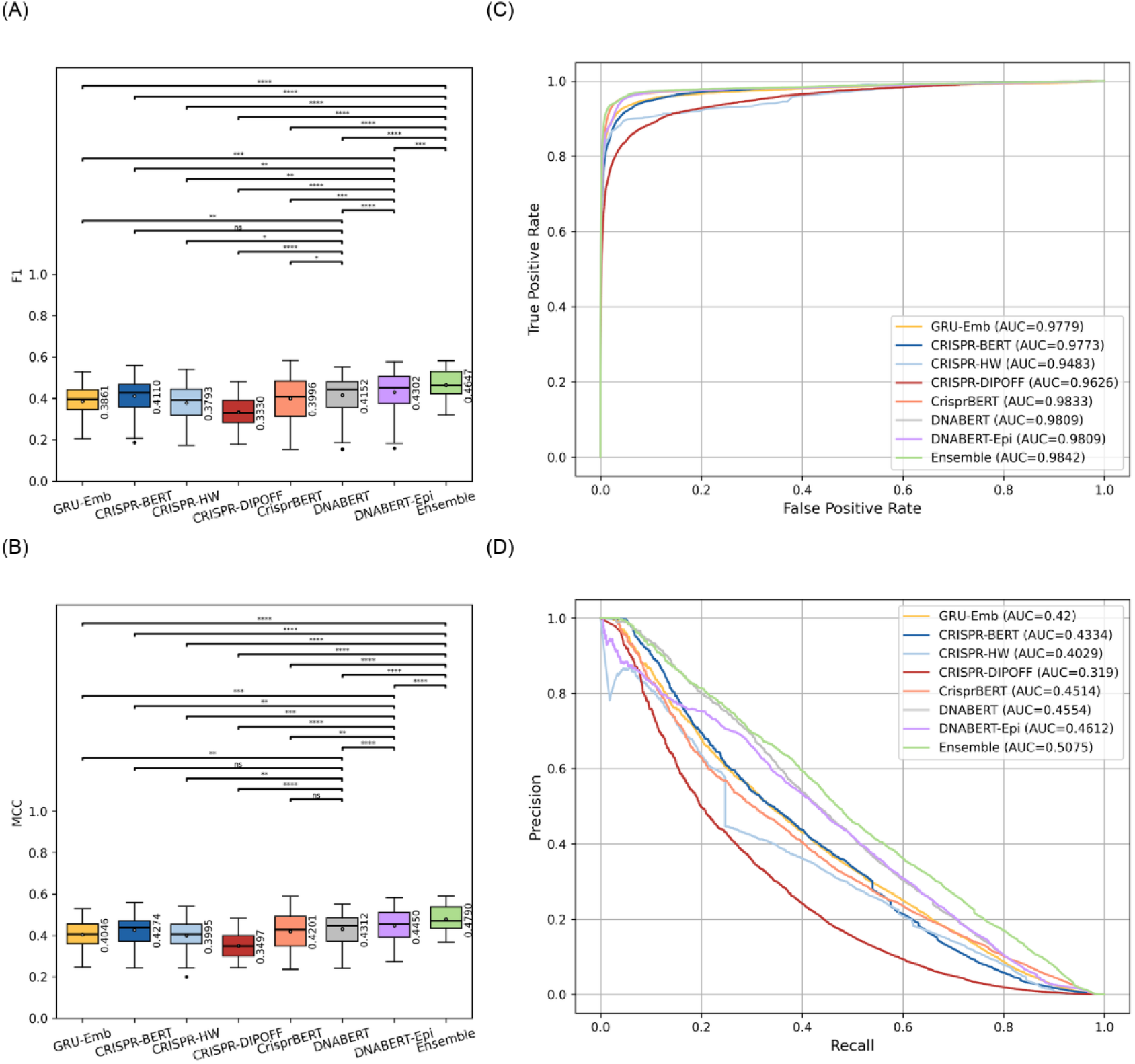
Performance comparison of models trained on the CHANGE-seq dataset and fine-tuned and tested on the GUIDE-seq dataset. (A) and (B) show boxplots of the F1 score and Matthews correlation coefficient (MCC), respectively, for each model. Statistical significance between models is indicated by asterisks (ns: no significance, *: p < 0.05, **: p < 0.01, ***: p < 0.001, ****: p < 0.0001). (C) presents the receiver operating characteristic (ROC) curves, while (D) shows the precision-recall (PR) curves, both generated using all predicted probabilities from the experiment. The area under the curve (AUC) values in the legends represent the mean AUC scores obtained from cross-validation.

**Fig 4.**
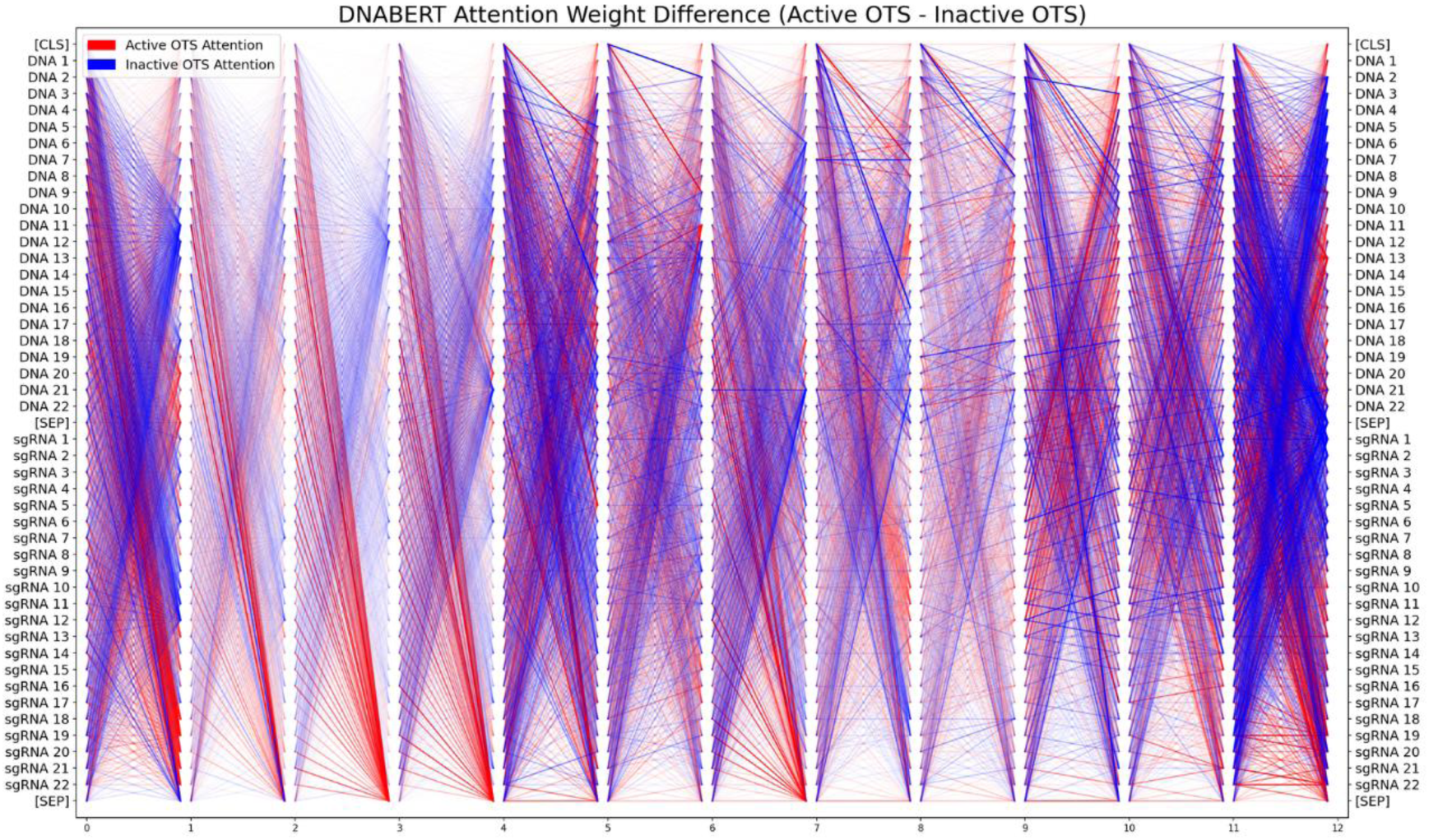
Visualization of attention weights differences between active and inactive OTS across DNABERT’s 12 transformer layers. The vertical axis represents input sequence tokens, while the horizontal axis represents transformer layers. Red lines indicate higher attention in active OTS, and blue lines indicate higher attention in inactive OTS.

The most significant differences were observed in the final transformer layer, where the active OTS exhibited a stronger focus on the latter half of the sequence (positions 10’–22’), whereas the inactive OTS emphasized the first half (positions 1’–10’). This pattern was consistent across multiple DNA-sgRNA sequences, suggesting that DNABERT prioritizes specific sequence regions when predicting off-target activity.

## Discussion

This study has made several contributions to the field of CRISPR/Cas9 off-target prediction. First, we applied the pre-trained DNABERT model to the off-target prediction task for the first time and compared its performance with state-of-the-art baseline models. Additionally, we conducted an ablation study on the effect of pre-training, demonstrating that masked language model (MLM) pre-training is essential for fine-tuning DNABERT on the off-target prediction task. This can be attributed to the large number of parameters in DNABERT, which makes optimization from random initialization highly difficult, especially in the context of imbalanced and noisy biological data. Without pre-training, the model fails to capture meaningful sequence patterns, as evidenced by stagnant loss values and near-zero evaluation metrics. In contrast, MLM pre-training enables DNABERT to acquire generalizable representations of DNA sequences, including k-mer dependencies and biologically relevant motifs, which facilitate more effective fine-tuning. Such prior knowledge helps the model to better distinguish subtle mismatches and sequence contexts that are critical for predicting off-target effects. Second, we proposed DNABERT-Epi, a method that integrates DNABERT with epigenetic features. We showed that DNABERT-Epi outperforms baseline models in off-target effect prediction and, through ablation study, demonstrated that epigenetic features significantly enhance predictive performance. Third, we demonstrated that an ensemble strategy combining multiple models substantially improves prediction performance. Individual models, including DNABERT-Epi and baseline models, performed worse than the ensemble model across all evaluation metrics.

This suggests that an ensemble approach could be practically useful for CRISPR/Cas9 experiments. Fourth, we attempted to interpret DNABERT’s internal processing by visualizing attention weights in the transformer layers. Notably, we identified distinct patterns in the attention weights of the final transformer layer. This study revealed that ensemble modeling significantly improves prediction performance, a phenomenon also observed in previous research [39]. Interestingly, we found that removing any single model from the ensemble led to a decline in performance. For example, in Scenario 3, excluding CRISPR-DIPOFF, the baseline model with the lowest performance, resulted in lower scores for F1, MCC, ROC-AUC, and PR-AUC (0.4577±0.0793, 0.4717±0.0671, 0.9836±0.0107, 0.4994±0.0633, respectively). This suggests that CRISPR-DIPOFF captures unique features that play an essential role in the ensemble, and its exclusion caused a greater performance decrease than when all models were included in the ensemble. A similar trend was observed when other models were excluded from the ensemble, further emphasizing the importance of each model’s contribution. This reinforces the idea that ensemble models benefit from the complementary strengths of individual models.

Our visualization of DNABERT’s attention weights revealed distinctive patterns, particularly in the final layer. Active OTS received higher attention in the latter part of the sequence, whereas inactive OTS were more focused on the earlier regions. This implies that DNABERT differentiates off-target effects based on these patterns. In CRISPR/Cas9 systems, the sgRNA and target DNA sequences contain a “seed region,” which is critical for target recognition [7, 40–45]. Cas9 requires perfect complementarity in this seed region, while it tolerates mismatches in the non-seed region. Our findings suggest that DNABERT learns this pattern from the dataset and prioritizes the seed region when predicting off-target effects.

However, several challenges remain. Although DNABERT-Epi outperformed the baseline models, the improvement was not substantial. This indicates that the epigenetic information used in this study (H3K4me3, ATAC, H3K27ac) and the model’s capacity may be insufficient to fully capture off-target effects. Future research should consider incorporating additional biological factors, such as the three-dimensional structure of target DNA and the secondary structure of sgRNA. Another challenge is the decline in predictive accuracy when the number of mismatches between sgRNA and DNA is high (5-6 mismatches; see S7-S9 Table). In such cases, active OTS were frequently misclassified as inactive. This issue was common across all models used in this study and may stem from the models’ inability to learn the binding and cleavage characteristics of Cas9 when multiple mismatches are present. To address this, future research should explore specialized modeling techniques for high-mismatch cases. Furthermore, although we visualized DNABERT’s attention weights, fully interpreting its internal mechanisms remains an open question. While we focused on the final transformer’s attention weights and their relationship with the seed region, the biological significance of attention patterns in intermediate layers remains unclear. Future studies should leverage additional interpretability methods, such as SHAP values and Grad-CAM, to gain deeper insights into the model’s decision-making process.

In summary, further advancements in CRISPR/Cas9 off-target prediction require integrating diverse biological features, improving model performance for challenging cases, and enhancing interpretability. These efforts will contribute to improving the accuracy of off-target predictions and increasing the practical applicability of CRISPR/Cas9 in experimental settings.

## Conclusion

Our study presents the first application of a pre-trained transformer-based model to CRISPR/Cas9 off-target prediction, demonstrating that DNABERT-Epi effectively integrates sequence and epigenetic features for improved accuracy. Our results suggest that leveraging both pretraining and multi-modal genomic information is key to enhancing predictive performance. Future work should focus on refining interpretability methods and expanding validation datasets to further advance computational CRISPR off-target prediction.

## Acknowledgements

In this research, the super-computing resource was provided by Human Genome Center, the Institute of Medical Science, the University of Tokyo. In addition, computations were partially performed on the NIG supercomputer at ROIS National Institute of Genetics.

## Supporting information

**S1 Fig.**
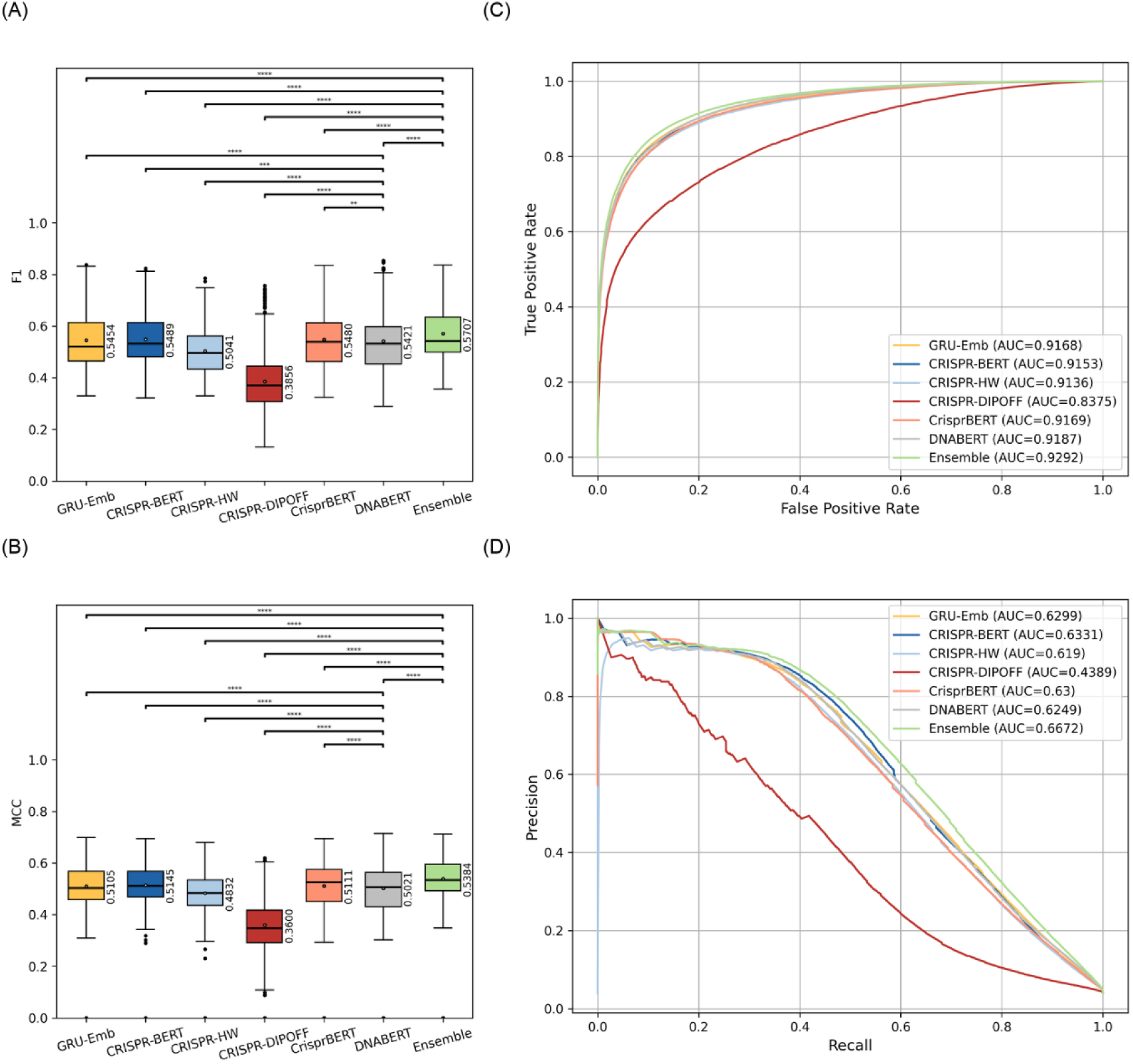
Performance comparison of models trained and tested on the CHANGE-seq dataset for four key metrics (F1, MCC, ROC-AUC, and PR-AUC). (A) and (B) show boxplots of the F1 score and Matthews correlation coefficient (MCC), respectively, for each model. Statistical significance between models is indicated by asterisks (ns: no significance, *: p < 0.05, **: p < 0.01, ***: p < 0.001, ****: p < 0.0001). (C) presents the receiver operating characteristic (ROC) curves, while (D) shows the precision-recall (PR) curves, both generated using all predicted probabilities from the experiment. The area under the curve (AUC) values in the legends represent the mean AUC scores obtained from cross-validation.

**S2 Fig.**
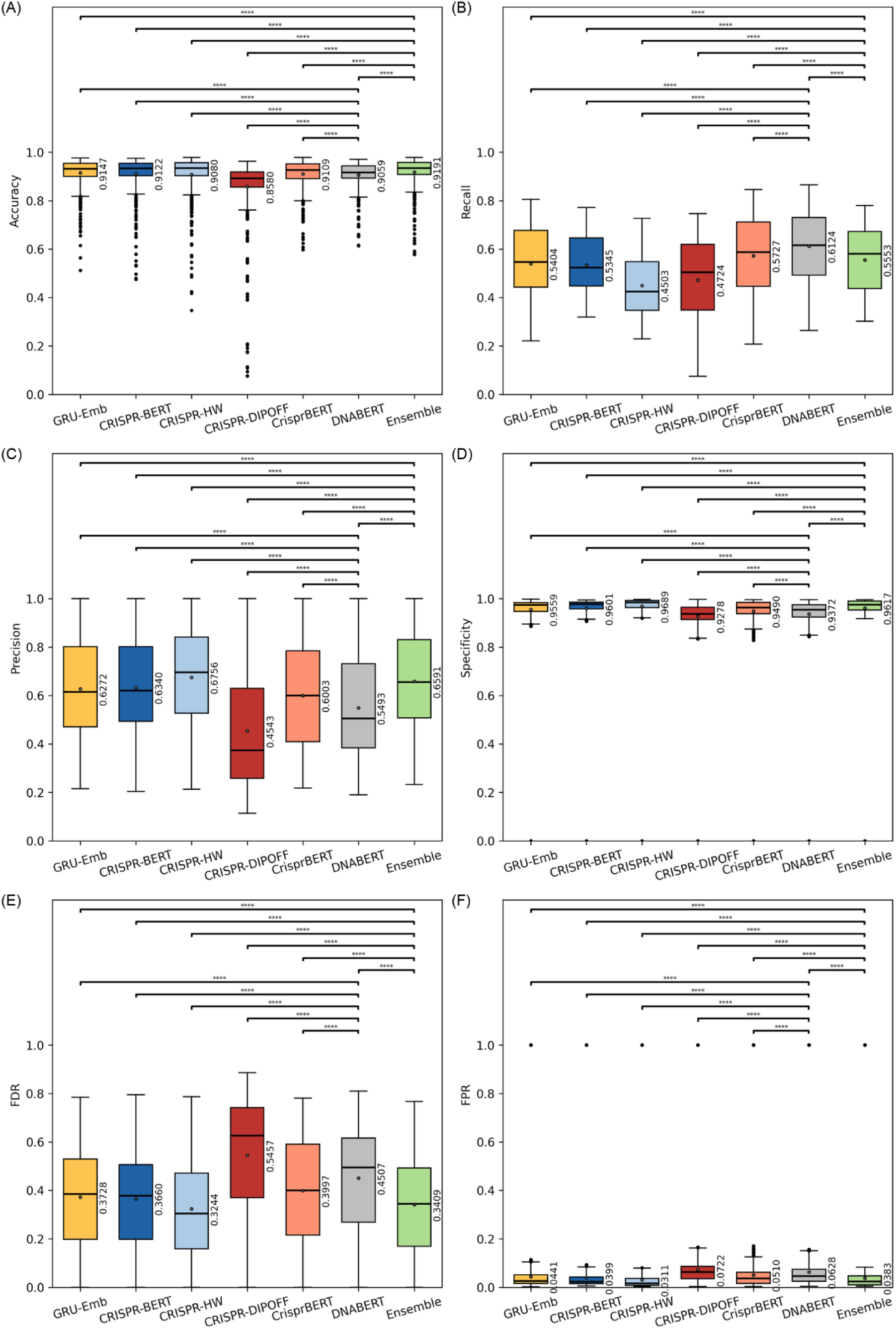
Performance comparison of models trained and tested on the CHANGE-seq dataset for six additional metrics (Accuracy, Recall, Precision, Specificity, FDR, and FPR). (A)–(F) show boxplots of Accuracy, Recall, Precision, Specificity, false discovery rate (FDR), and false positive rate (FPR) for each model. Statistical significance between models is indicated by asterisks based on the Wilcoxon test (ns: no significance, *: p < 0.05, **: p < 0.01, ***: p < 0.001, ****: p < 0.0001).

**S3 Fig.**
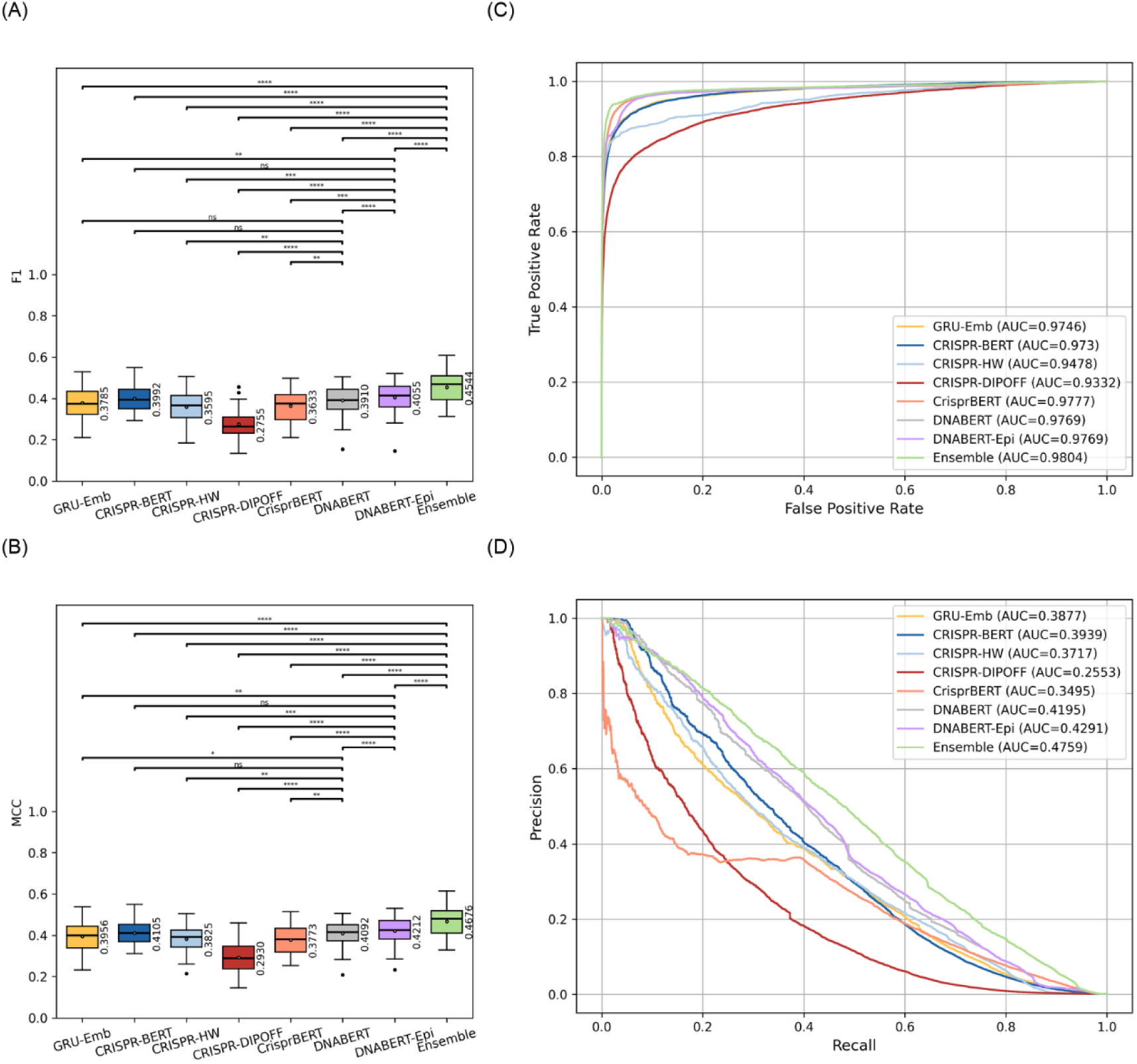
Performance comparison of models trained and tested on the GUIDE-seq dataset for four key metrics (F1, MCC, ROC-AUC, and PR-AUC). (A) and (B) show boxplots of the F1 score and Matthews correlation coefficient (MCC), respectively, for each model. Statistical significance between models is indicated by asterisks (ns: no significance, *: p < 0.05, **: p < 0.01, ***: p < 0.001, ****: p < 0.0001). (C) presents the receiver operating characteristic (ROC) curves, while (D) shows the precision-recall (PR) curves, both generated using all predicted probabilities from the experiment. The area under the curve (AUC) values in the legends represent the mean AUC scores obtained from cross-validation.

**S4 Fig.**
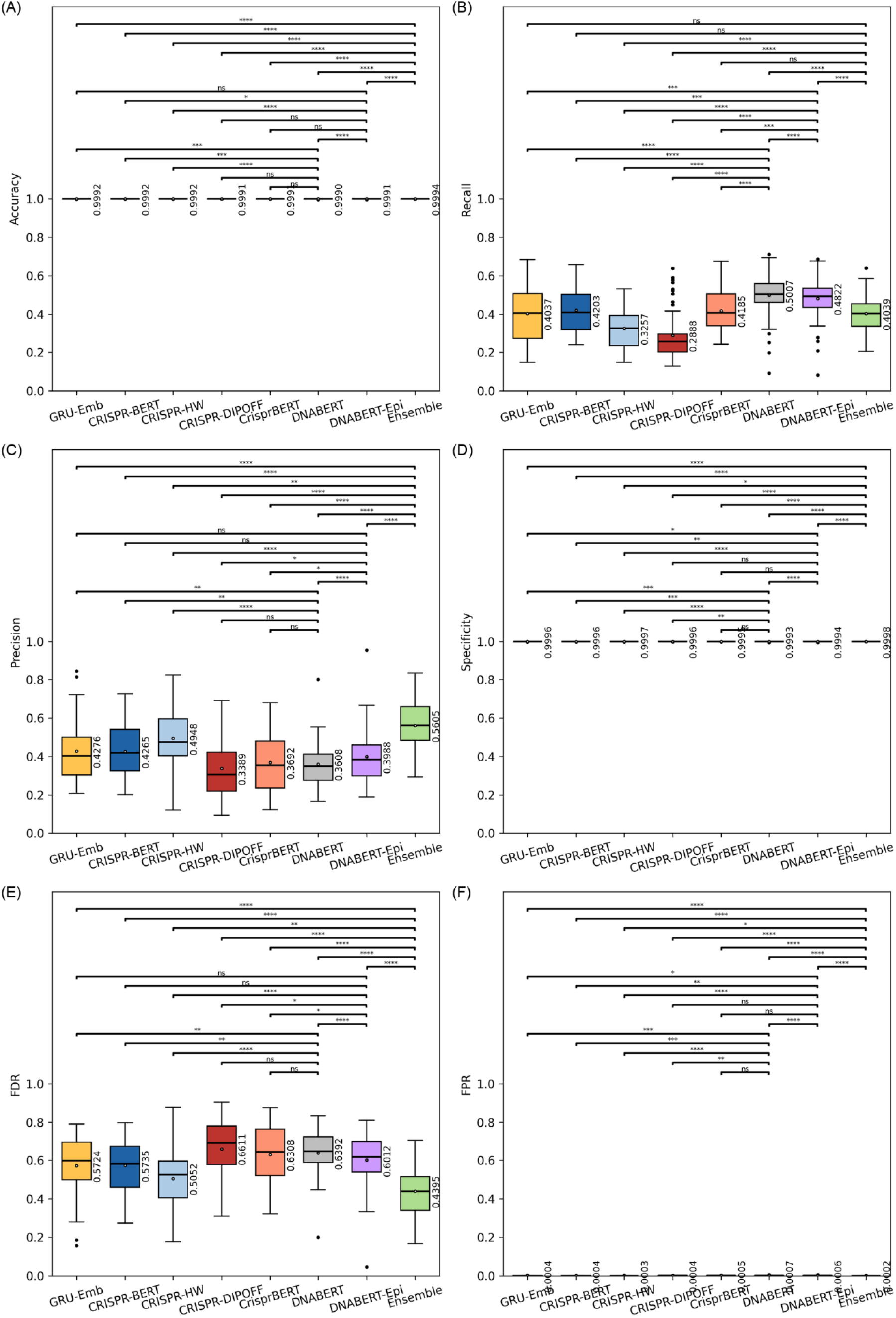
Performance comparison of models trained and tested on the GUIDE-seq dataset for six additional metrics (Accuracy, Recall, Precision, Specificity, FDR, and FPR). (A)–(F) show boxplots of Accuracy, Recall, Precision, Specificity, false discovery rate (FDR), and false positive rate (FPR) for each model. Statistical significance between models is indicated by asterisks based on the Wilcoxon test (ns: no significance, *: p < 0.05, **: p < 0.01, ***: p < 0.001, ****: p < 0.0001).

**S5 Fig.**
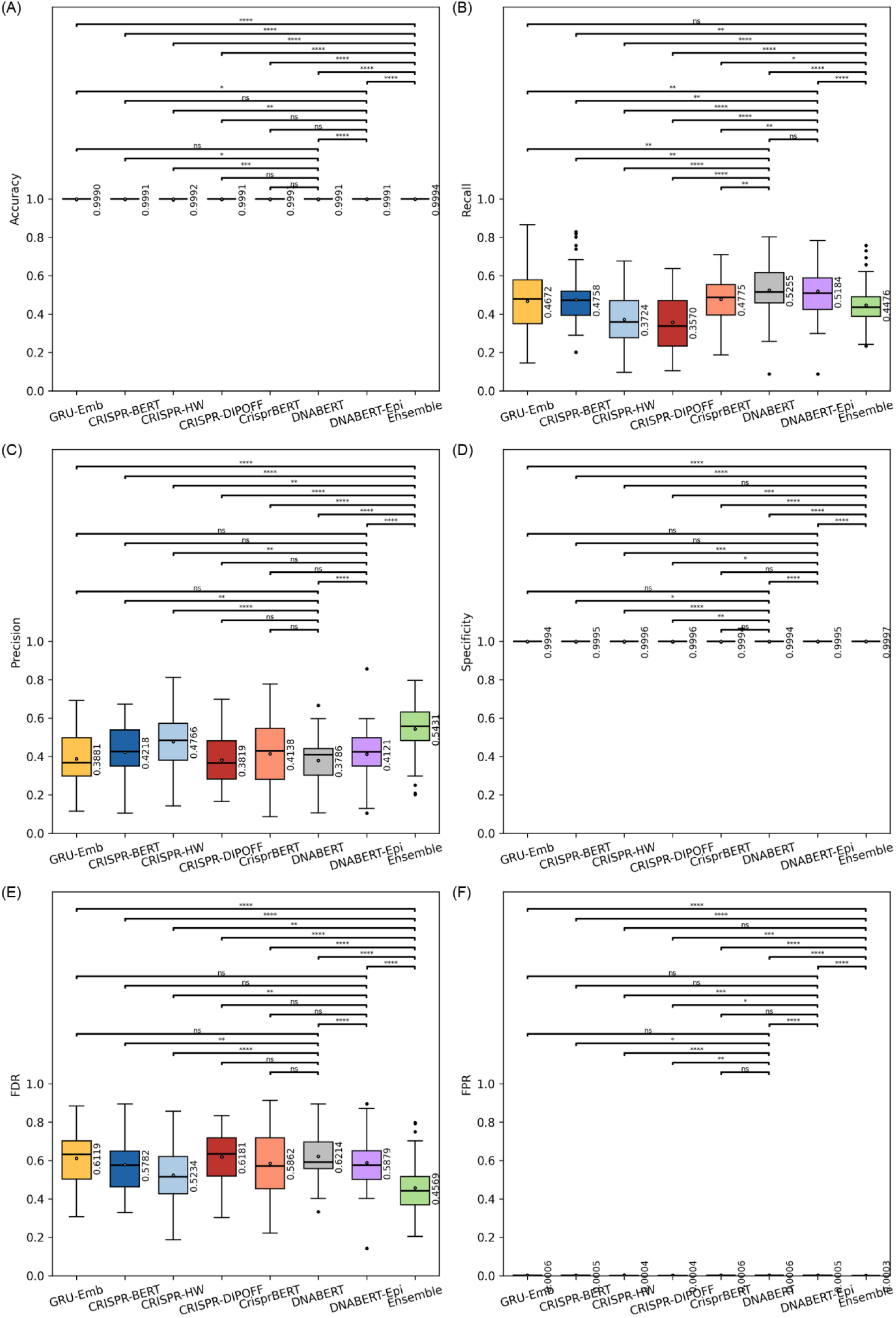
Performance comparison of models trained on the CHANGE-seq dataset and fine-tuned and tested on the GUIDE-seq dataset for six additional metrics (Accuracy, Recall, Precision, Specificity, FDR, and FPR). (A)–(F) show boxplots of Accuracy, Recall, Precision, Specificity, false discovery rate (FDR), and false positive rate (FPR) for each model. Statistical significance between models is indicated by asterisks based on the Wilcoxon test (ns: no significance, *: p < 0.05, **: p < 0.01, ***: p < 0.001, ****: p < 0.0001).

**S6 Fig.**
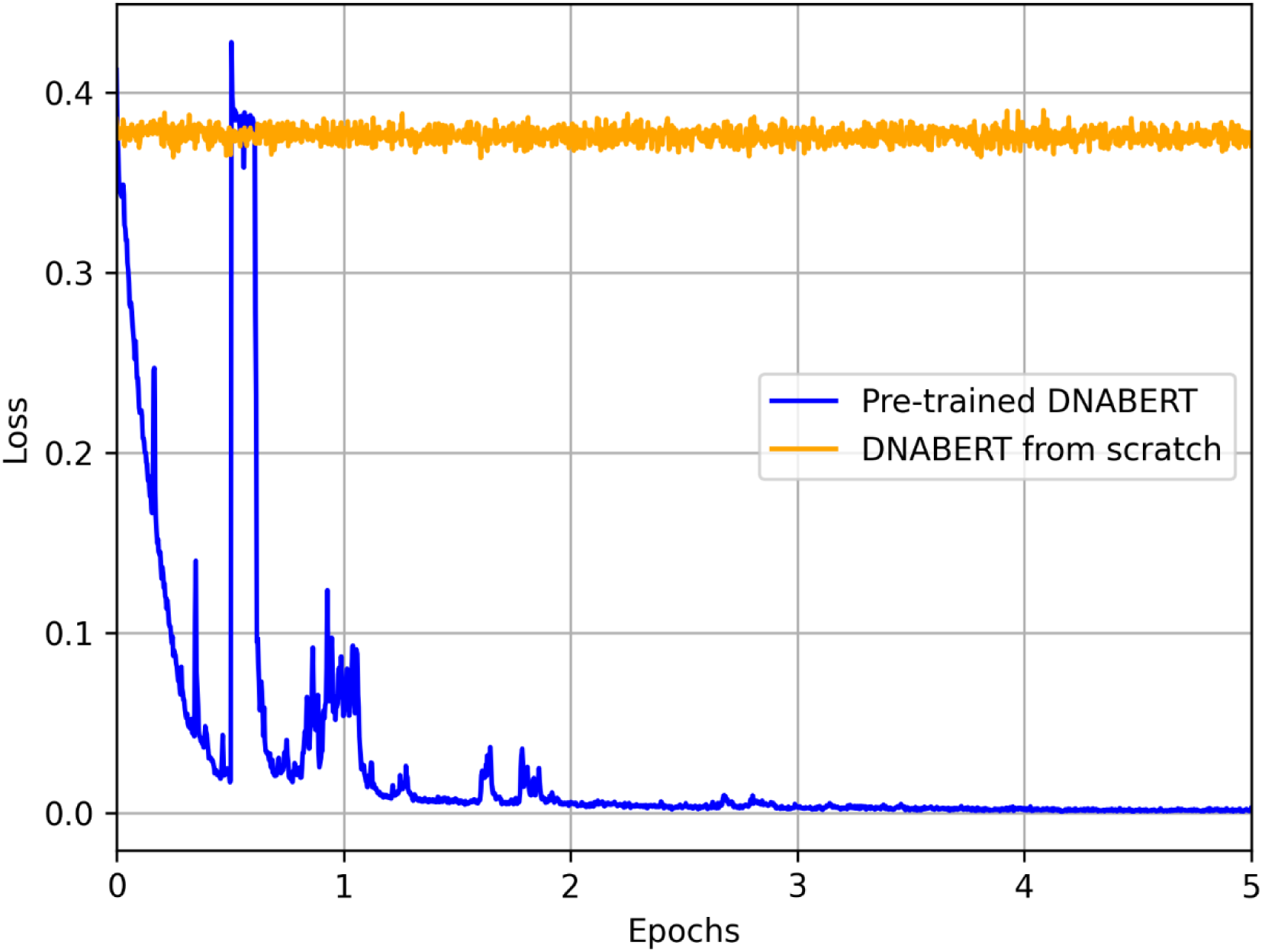
Training loss curves during mismatch prediction fine-tuning for pre-trained and randomly initialized DNABERT. Training loss curves for the mismatch prediction task during fine-tuning. The pre-trained DNABERT (blue line) exhibited a consistent decrease in loss as training progressed, indicating effective learning. In contrast, DNABERT from scratch (orange line) showed a flat loss curve, suggesting a lack of meaningful pattern recognition. The x-axis represents training progress in epochs, while the y-axis denotes the loss value.

**S1 Table. Performance comparison of models trained and tested on the CHANGE-seq dataset.** This table compares the performance of each model using various evaluation metrics, including accuracy, recall, precision, specificity, false positive rate (FPR), false discovery rate (FDR), F1 score, ROC-AUC (receiver operating characteristic area under the curve), PR-AUC (precision-recall area under the curve), and MCC (Matthews correlation coefficient). For each metric, the table provides a detailed statistical breakdown, including the mean, median, standard deviation (std), maximum (max), and minimum (min) values.

**S2 Table. Wilcoxon signed-rank test results for models trained and tested on the CHANGE-seq dataset.** This table presents the p-values from the Wilcoxon signed-rank test, used to evaluate performance differences between all model pairs. The performance metrics analyzed include accuracy, recall, precision, specificity, false positive rate (FPR), false discovery rate (FDR), F1 score, ROC-AUC (receiver operating characteristic area under the curve), PR-AUC (precision-recall area under the curve), and MCC (Matthews correlation coefficient).

**S3 Table. Performance comparison of models trained and tested on the GUIDE-seq dataset.** This table compares the performance of each model using evaluation metrics such as accuracy, recall, precision, specificity, false positive rate (FPR), false discovery rate (FDR), F1 score, ROC-AUC, PR-AUC, and MCC. Each metric includes a detailed statistical summary comprising the mean, median, standard deviation (std), maximum (max), and minimum (min) values.

**S4 Table. Wilcoxon signed-rank test results for models trained and tested on the GUIDE-seq dataset.** This table shows the p-values from the Wilcoxon signed-rank test conducted to compare the performance of different models. The metrics evaluated include accuracy, recall, precision, specificity, false positive rate (FPR), false discovery rate (FDR), F1 score, ROC-AUC, PR-AUC, and MCC.

**S5 Table. Performance comparison of models trained on the CHANGE-seq dataset and fine-tuned and tested on the GUIDE-seq.** This table compares the performance of each model using evaluation metrics such as accuracy, recall, precision, specificity, false positive rate (FPR), false discovery rate (FDR), F1 score, ROC-AUC, PR-AUC, and MCC. Each metric includes a detailed statistical summary comprising the mean, median, standard deviation (std), maximum (max), and minimum (min) values.

**S6 Table. Wilcoxon signed-rank test results for models trained on the CHANGE-seq dataset and fine-tuned and tested on the GUIDE-seq.** This table shows the p-values from the Wilcoxon signed-rank test conducted to compare the performance of different models. The metrics evaluated include accuracy, recall, precision, specificity, false positive rate (FPR), false discovery rate (FDR), F1 score, ROC-AUC, PR-AUC, and MCC.

**S7 Table. Confusion matrices of models trained and tested on the CHANGE-seq dataset.** This table presents the confusion matrices for each model. The values in each matrix represent the aggregated results from five independent experimental runs.

**S8 Table. Confusion matrices of models trained and tested on the GUIDE-seq dataset.** This table presents the confusion matrices for each model. The values in each matrix represent the aggregated results from five independent experimental runs.

**S9 Table. Confusion matrices of models trained on the CHANGE-seq dataset and fine-tuned and tested on the GUIDE-seq.** This table presents the confusion matrices for each model. The values in each matrix represent the aggregated results from five independent experimental runs.

## References

1. Jinek M, Chylinski K, Fonfara I, Hauer M, Doudna JA, Charpentier E. A programmable dual-rna–guided dna endonuclease in adaptive bacterial immunity. Science. 2012; 337(6096): 816–821. doi: 10.1126/science.1225829

2. Doudna JA, Charpentier E. The new frontier of genome engineering with CRISPR-Cas9. Science. 2014; 346(6213): 1258096. doi: 10.1126/science.1258096

3. Sharma G, Sharma AR, Bhattacharya M, Lee SS. CRISPR-Cas9: A Preclinical and Clinical Perspective for the Treatment of Human Diseases. Mol Ther. 2021; 29(2): 571–586. doi: 10.1016/j.ymthe.2020.09.028

4. Hsu PD, Lander ES, Zhang F. Development and applications of crispr-cas9 for genome engineering. Cell. 2014; 157(6): 1262–1278. doi: 10.1016/j.cell.2014.05.010

5. Deveau H, Garneau JE, Moineau S. CRISPR/Cas system and its role in phage-bacteria interactions. Annu. Rev. Microbiol. 2010; 64: 475–493. doi: 10.1146/annurev.micro.112408.134123

6. Horvath P, Barrangou R. CRISPR/Cas, the Immune System of Bacteria and Archaea. Science. 2010; 327(5962): 167–170. doi: 10.1126/science.1179555

7. Zhang XH, Tee LY, Wang XG, Huang QS, Yang SH. Off-target Effects in CRISPR/Cas9-mediated Genome Engineering. Mol Ther Nucleic Acids. 2015; 4: e264. doi: 10.1038/mtna.2015.37

8. Mali P, Aach J, Stranges PB, Esvelt KM, Moosburner M, Kosuri S, et al. CAS9 transcriptional activators for target specificity screening and paired nickases for cooperative genome engineering. Nat Biotechnol. 2013; 31(9): 833–838. doi: 10.1038/nbt.2675

9. Chen JS, Dagdas YS, Kleinstiver BP, Welch MM, Sousa AA, Harrington LB, et al. Enhanced proofreading governs CRISPR-Cas9 targeting accuracy. Nature. 2017; 550(7676): 407–410. doi: 10.1038/nature24268

10. Lin J, Wong KC. Off-target predictions in CRISPR-Cas9 gene editing using deep learning. Bioinformatics. 2018; 34(17): i656–i663. doi: 10.1093/bioinformatics/bty554

11. Manghwar H, Li B, Ding X, Hussain A, Lindsey K, Zhang X, et al. CRISPR/Cas Systems in Genome Editing: Methodologies and Tools for sgRNA Design, Off-Target Evaluation, and Strategies to Mitigate Off-Target Effects. Adv. Sci. 2020; 7(6): 1902312. doi: 10.1002/advs.201902312.

12. Peng R, Lin G, Li J. Potential pitfalls of CRISPR/Cas9-mediated genome editing. FEBS J. 2016; 283(7): 1218–1231. doi: 10.1111/febs.13586

13. Sherkatghanad Z, Abdar M, Charlier J, Makarenkov V. Using traditional machine learning and deep learning methods for on- and off-target prediction in CRISPR/Cas9: a review. Brief Bioinform. 2023; 24(3): bbad131. doi: 10.1093/bib/bbad131

14. Konstantakos V, Nentidis A, Krithara A, Paliouras G. CRISPR–Cas9 gRNA efficiency prediction: an overview of predictive tools and the role of deep learning. Nucleic Acids Res. 2022; 50(7): 3616–3637. doi: 10.1093/nar/gkac192

15. Zhang G, Luo Y, Dai X, Dai Z. Benchmarking deep learning methods for predicting CRISPR/Cas9 sgRNA on- and off-target activities. Brief Bioinform. 2023; 24(6): bbad333. doi: 10.1093/bib/bbad333

16. Zhang ZR, Jiang ZR. Effective use of sequence information to predict CRISPR-Cas9 off-target. Comput Struct Biotechnol J. 2022; 20: 650–661. doi: 10.1016/j.csbj.2022.01.006

17. Liu Q, He D, Xie L. Prediction of off-target specificity and cell-specific fitness of CRISPR-Cas System using attention boosted deep learning and network-based gene feature. PLoS Comput Biol. 2019; 15(10): e1007480. doi: 10.1371/journal.pcbi.1007480

18. Zhang G, Zeng T, Dai Z, Dai X. Prediction of CRISPR/Cas9 single guide RNA cleavage efficiency and specificity by attention-based convolutional neural networks. Comput Struct Biotechnol J. 2021; 19: 1445–1457. doi: 10.1016/j.csbj.2021.03.001

19. Guan Z, Jiang Z. Transformer-based anti-noise models for CRISPR-Cas9 off-target activities prediction. Brief Bioinform. 2023; 24(3): bbad127. doi: 10.1093/bib/bbad127

20. Sun J, Guo J, Liu J. CRISPR-M: Predicting sgRNA off-target effect using a multi-view deep learning network. PLoS Comput Biol. 2024; 20(3): e1011972. doi: 10.1371/journal.pcbi.1011972

21. Chen D, Shu W, Peng S. Predicting CRISPR-Cas9 Off-target with Self-supervised Neural Networks. In: Proceedings of the 2020 IEEE International Conference on Bioinformatics and Biomedicine (BIBM); 2020 Dec 9-12; Washington, DC, USA. IEEE; 2020. p.245–250. doi: 10.1109/BIBM49941.2020.9313280

22. Luo Y, Chen Y, Xie HZ, Zhu W, Zhang G. Interpretable CRISPR/Cas9 off-target activities with mismatches and indels prediction using BERT. Comput Biol Med. 2024; 169: 107932. doi: 10.1016/j.compbiomed.2024.107932

23. Sari O, Liu Z, Pan Y, Shao X. Predicting CRISPR-Cas9 off-target effects in human primary cells using bidirectional LSTM with BERT embedding. Bioinformatics Adv. 2025; 5(1): vbae184. doi: 10.1093/bioadv/vbae184

24. Lazzarotto CR, Malinin NL, Li Y, Zhang R, Yang Y, Lee G, et al. CHANGE-seq reveals genetic and epigenetic effects on CRISPR–Cas9 genome-wide activity. Nat Biotechnol. 2020; 38(11): 1317–1327. doi: 10.1038/s41587-020-0555-7

25. Chuai G, Ma H, Yan J, Chen M, Hong N, Xue D, et al. DeepCRISPR: optimized CRISPR guide RNA design by deep learning. Genome Biol. 2018; 19(1): 80. doi: 10.1186/s13059-018-1459-4

26. Zhang G, Dai Z, Dai X. C-RNNCrispr: Prediction of CRISPR/Cas9 sgRNA activity using convolutional and recurrent neural networks. Comput Struct Biotechnol J. 2020; 18: 344–354. doi: 10.1016/j.csbj.2020.01.013

27. Mak JK, Störtz F, Minary P. Comprehensive computational analysis of epigenetic descriptors affecting CRISPR-Cas9 off-target activity. BMC Genomics. 2022; 23(1): 805. doi: 10.1186/s12864-022-09012-7

28. Ji Y, Zhou Z, Liu H, Davuluri RV. DNABERT: pre-trained Bidirectional Encoder Representations from Transformers model for DNA-language in genome. Bioinformatics. 2021; 37(15): 2112–2120. doi: 10.1093/bioinformatics/btab083

29. Tsai SQ, Zheng Z, Nguyen NT, Liebers M, Topkar VV, Thapar V, et al. GUIDE-seq enables genome-wide profiling of off-target cleavage by CRISPR-Cas nucleases. Nat Biotechnol. 2015; 33(2): 187–197. doi: 10.1038/nbt.3117

30. Yaish O, Orenstein Y. Generating, modeling and evaluating a large-scale set of CRISPR/Cas9 off-target sites with bulges. Nucleic Acids Res. 2024; 52(12): 6777–6790. doi: 10.1093/nar/gkae428

31. Akcakaya P, Bobbin ML, Guo JA, Malagon-Lopez J, Clement K, Garcia SP, et al. In vivo CRISPR editing with no detectable genome-wide off-target mutations. Nature. 2018; 561(7723): 416-419. doi: 10.1038/s41586-018-0500-9

32. Tsai SQ, Nguyen NT, Malagon-Lopez J, Topkar VV, Aryee MJ, Joung JK. CIRCLE-seq: a highly sensitive in vitro screen for genome-wide CRISPR–Cas9 nuclease off-targets. Nat Methods. 2017; 14(6): 607–614. doi: 10.1038/nmeth.4278

33. Chung CH, Allen AG, Sullivan NT, Atkins A, Nonnemacher MR, Wigdahl B, et al. Computational Analysis Concerning the Impact of DNA Accessibility on CRISPR-Cas9 Cleavage Efficiency. Mol Ther. 2020; 28(1): 19–28. doi: 10.1016/j.ymthe.2019.10.008

34. Yaish O, Asif M, Orenstein Y. A systematic evaluation of data processing and problem formulation of CRISPR off-target site prediction. Brief Bioinform. 2022; 23(5): bbac157. doi: 10.1093/bib/bbac157

35. Yang Y, Li J, Zou Q, Ruan Y, Feng H. Prediction of CRISPR-Cas9 off-target activities with mismatches and indels based on hybrid neural network. Comput Struct Biotechnol J. 2023; 21: 5039–5048. doi: 10.1016/j.csbj.2023.10.018

36. Toufikuzzaman M, Hassan Samee MA, Sohel Rahman M. CRISPR-DIPOFF: an interpretable deep learning approach for CRISPR Cas-9 off-target prediction. Brief Bioinform. 2024; 25(2): bbad530. doi: 10.1093/bib/bbad530

37. Vig J. A Multiscale Visualization of Attention in the Transformer Model. arXiv. 2019. doi: 10.48550/arXiv.1906.05714

38. Wang ZJ, Turko R, Chau DH. Dodrio: Exploring Transformer Models with Interactive Visualization. arXiv. 2021.doi: 10.48550/arXiv.2103.14625

39. Zhang S, Li X, Lin Q, Wong K-C. Synergizing CRISPR/Cas9 off-target predictions for ensemble insights and practical applications. Bioinformatics. 2019; 35(7): 1108–1115. doi: 10.1093/bioinformatics/bty748

40. Pacesa M, Lin C-H, Cléry A, Saha A, Arantes PR, Bargsten K, et al. Structural basis for Cas9 off-target activity. Cell. 2022; 185(22): 4067–4081.e21. doi: 10.1016/j.cell.2022.09.026

41. Semenova E, Jore MM, Datsenko KA, Semenova A, Westra ER, Wanner B, et al. Interference by clustered regularly interspaced short palindromic repeat (CRISPR) RNA is governed by a seed sequence. Proc Natl Acad Sci U S A. 2011; 108(25): 10098–10103. doi: 10.1073/pnas.1104144108

42. Jiang F, Doudna JA. CRISPR–Cas9 Structures and Mechanisms. Annu Rev Biophys. 2017; 46: 505–529. doi: 10.1146/annurev-biophys-062215-010822

43. Yu T, Liu T, Wang Y, Zhao X, Zhang W. Effect of Cas9 Protein on the Seed-Target Base Pair of the sgRNA/DNA Hybrid Duplex. J Phys Chem B. 2023; 127(22): 4989–4997. doi: 10.1021/acs.jpcb.3c00997

44. Feng Y, Liu S, Chen R, Xie A. Target binding and residence: a new determinant of DNA double-strand break repair pathway choice in CRISPR/Cas9 genome editing. J. Zhejiang Univ. Sci. B. 2021; 22: 73–86. doi: 10.1631/jzus.B2000282

45. Hsu PD, Scott DA, Weinstein JA, Ran FA, Konermann S, Agarwala V, et al. DNA targeting specificity of RNA-guided Cas9 nucleases. Nat Biotechnol. 2013; 31(9): 827–832. doi: 10.1038/nbt.2647

